# Accessible and interactive RNA sequencing analysis using serverless computing

**DOI:** 10.1101/576199

**Authors:** Ling-Hong Hung, Xingzhi Niu, Wes Lloyd, Ka Yee Yeung

## Abstract

We present a novel method that yields 1100-fold computational speedup and allows biomedical scientists to interactively adjust alignment parameters in real time to iteratively improve final analytical results. Specifically, the alignment time for a 640 million read human RNA-sequencing dataset is reduced from 19 hours to 1 minute using serverless cloud computing. We provide a graphical interface for the accelerated workflows, thus making our methodology accessible to non-cloud experts.

---

For the analyses of RNA-sequencing (RNA-seq) data, the most computationally intensive step is the mapping of reads to the reference sequence. Although many optimized algorithms and software tools have been developed for this step^1–3^, the time required for alignment to a reference remains substantial. Consequently, once data is aligned, the analysis is rarely repeated even though parameter choices greatly influence the quality of the alignment^4^, potentially affecting downstream results and conclusions. We present a novel methodology using serverless cloud computing to ***reduce the execution time for alignment by 1100-fold***. We incorporate this methodology into a workflow that determines differentially expressed genes (DEGs). We provide a graphical interface that enables biomedical users who are not cloud experts to interactively tune alignment parameters. In a case study, we demonstrate that changing even a single alignment parameter can affect the composition of the list of top-10 DEGs.

To make interactive tuning of alignment parameters practical, the execution time required to align a dataset must be significantly reduced. Cloud computing has been used to accelerate the alignment process by using many virtual severs to simultaneously process multiple datasets^7–9^. While this approach is effective for analysis of large collections of samples, it does not improve the execution time for aligning an individual dataset. Other approaches that divide a single dataset into small pieces to be processed in parallel, and/or accelerate specific parts of the alignment algorithm, require specialized hardware^10^ or access to a private or cloud-based cluster^11^ for significant speedup. These solutions require significant expertise to set up and use. Recently, ***function-as-a-service (FaaS)*** cloud platforms provide on-demand access to small short-lived applications without the need to configure virtual servers before computation can proceed^5^. FaaS platforms provide ***serverless functions***, designed for small, short-lived microservices. These functions, however, have significant limitations with respect to memory, disk space, execution time, and network bandwidth. Using a variety of techniques, we overcome these limitations (see Methods) and harness ***serverless function instances*** as rapidly deployable, compute nodes to provide an on-demand cloud-based *“supercomputer”* for RNA-sequencing. Unlike approaches designed for processing large collections of samples^7–9^, our strategy supports accelerating the alignment of <u>a single dataset</u> on Amazon Web Services (AWS) or the Google Cloud Platform (GCP).

As a case study to demonstrate this novel use of serverless computing, we present an RNA-seq workflow using unique molecular identifiers (UMI) to obtain de-duped transcript counts^12^ which are then processed by edgeR^13^ to obtain a list of genes that were differentially expressed upon treatment with different combinations of drugs^14^. For the analysis of this dataset, we invoked 1752 function instances to **reduce the alignment time from 19.5 hours to 1 minute** when using the most performant configuration on AWS (Table S2). The dataset consists of 640 million reads (46 GB compressed files) obtained from cardiocytes treated with 15 different drug combinations for 48 hours on 96 well plates. The sequences contain a barcode sequence to identify the sample which allows for multiple samples to be pooled and loaded onto the same lane (multiplexed) for sequencing. The resulting fastq files of reads are processed using a 3-step pipeline to obtain the transcript counts. The first step uses the barcode sequences to separate or de-multiplex the reads into the 96 originating wells. These reads are then aligned to the human transcriptome using the Burrows-Wheeler Aligner (BWA)^1^. The resulting alignments are merged and de-duped using Unique Molecular Identifiers (UMIs) to compute the observed counts for each transcript. The original workflow^14^ took 29.5 hours to obtain transcript counts using a virtual cloud server (AWS EC2 instance m4.4xlarge). We previously reported optimizations of the implementation that reduce the total execution time to 3.5 hours when using 16 threads on the same type of cloud server^5^. As expected, the longest step is the CPU-intensive alignment which takes 19.5 hours in the original^14^ and 2.5 hours in the optimized implementation^5^. We benchmarked the serverless version of the workflow and observed that ***transcript counts can be obtained in 6 minutes at a cost of $3.85*** (see Methods for implementation details, benchmarking code and *cost calculations*). This execution time includes the time needed for all data transfers to the cloud and serverless instances and the total cost includes charges to rent the client computer. **Subsequent workflow executions take less than 2 minutes to obtain counts** as there is no need to repeat the generation and transfer of data shards.

We developed a containerized graphical front-end for the serverless version of the UMI RNA-seq workflow by extending our Biodepot-workflow-builder framework^14^ (see **Figure 1**), that can run from a laptop, desktop, or cloud server. The entire setup is automated with the user needing only to enter the credentials for the cloud account being used for the serverless instances, and the location of the files that are to be processed. The graphical tool uses the original edgeR script to obtain a final list of DEGs that is displayed in a fully functional spreadsheet. Our interface makes the workflow fully interactive, and the user can start, monitor, stop, modify, and re-start the analysis at any step with different parameter sets. The accessibility and minimal setup of the graphical workflow enables biomedical researchers who are not cloud experts to use a serverless supercomputer to rapidly modify alignment parameters and monitor the effect on the composition of the final gene lists.

**Figure 1.**
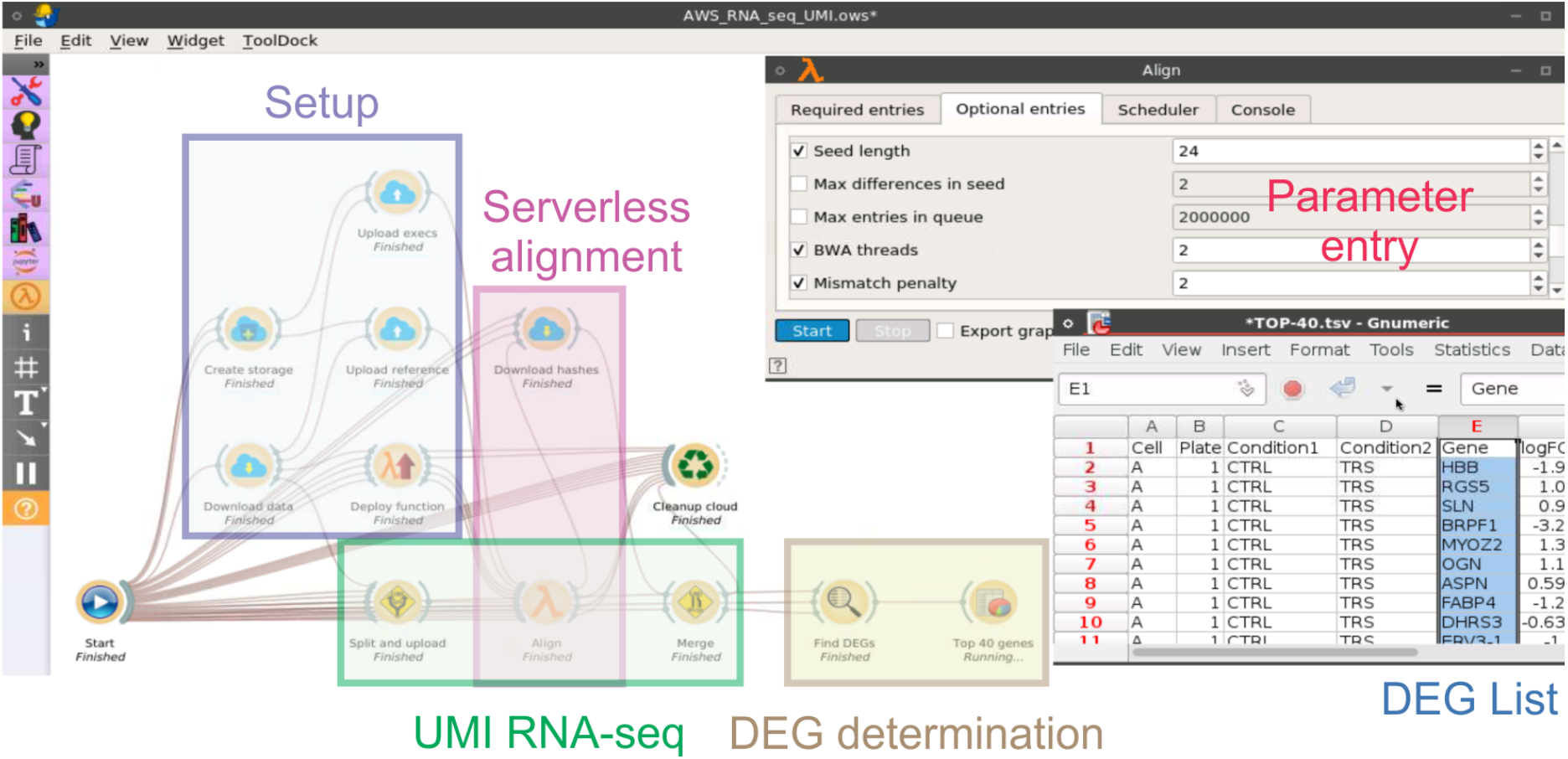
Graphical front-end for interactive analyses of differential expression using UMI RNA-seq and edgeR. A screenshot of the interactive front-end built with the Biodepot-workflow-builder, which can be run on a laptop, private, or cloud server and accessed using a browser. Each icon (widget) controls a separate containerized module. Double-clicking on a widget reveals graphical elements for parameter entry, starting and stopping execution, and displaying intermediate output. The parameter entry tab is shown for the Align widget in the upper left corner. Lines connecting widgets indicate data flow between the execution modules. Connections and widgets can be added and removed using a drag-and-drop interface. The workflow itself is started by double-clicking on the start widget in the lower left of the window. The user then enters the names of the data files and credentials directory into a form and presses a start button to begin execution. Execution proceeds automatically and finishes by popping up a fully functional spreadsheet populated with a list of DEGs. This is the window in the lower right. The user can monitor, stop, modify, and restart the workflow at any step. Colored boxes have been added to show the grouping of widgets involved in setup, alignment using serverless computing, the UMI RNA-seq pipeline, and DEG determination using edgeR. Other tools can alternatively be used for DEG. A cleanup module is also provided to delete intermediate results before re-running the workflow, or to delete all files and cloud resources after the workflow is complete. The front-end shown here depicts the Amazon Web Services (AWS) workflow. A GCP version is also available, and both are available from GitHub.

Default parameter settings for aligners are often not optimal^4^. Some parameter values are chosen to increase speed at the cost of accuracy. For example, BWA normally uses the leftmost base pairs of the sequence as a seed. Poor matches to the seed are immediately rejected, saving considerable time. Seedless alignment is available as an option which is more accurate but takes 2.5x longer^1^. This speed improvement becomes insignificant with the serverless implementation (Table 1C). Other parameters are tuned to specific properties of the data. For example, BWA sets the default maximum edit distance to 0.04 per 100 base pairs based on an expected number of sequence errors. This can be very different from the actual error rate. We showcase the flexibility of our tool to increase the maximum edit distance parameter, perform seedless alignment, and re-analyze the data from our case study in Table 1C. Our results illustrate how the overall transcript counts can be almost identical while producing changes in the top-10 DEG list. For transcripts with low counts, changes can cause them to become too low and be filtered by edgeR (Methods Figure S1). Changes in alignment parameters also alter the p-value and False Discovery Rate (FDR) which can affect the ordering and composition of DEG lists. In the top-10 list for alendronate treatment shown in Table 1A, we see that SLC90A4 drops out of the list and is replaced with SLC9A7 when we raise the maximum edit distance to 0.1 per 100 base pairs. This change in top-10 DEG list is confirmed using the more accurate seedless alignment in Table 1B.

**Table 1.**
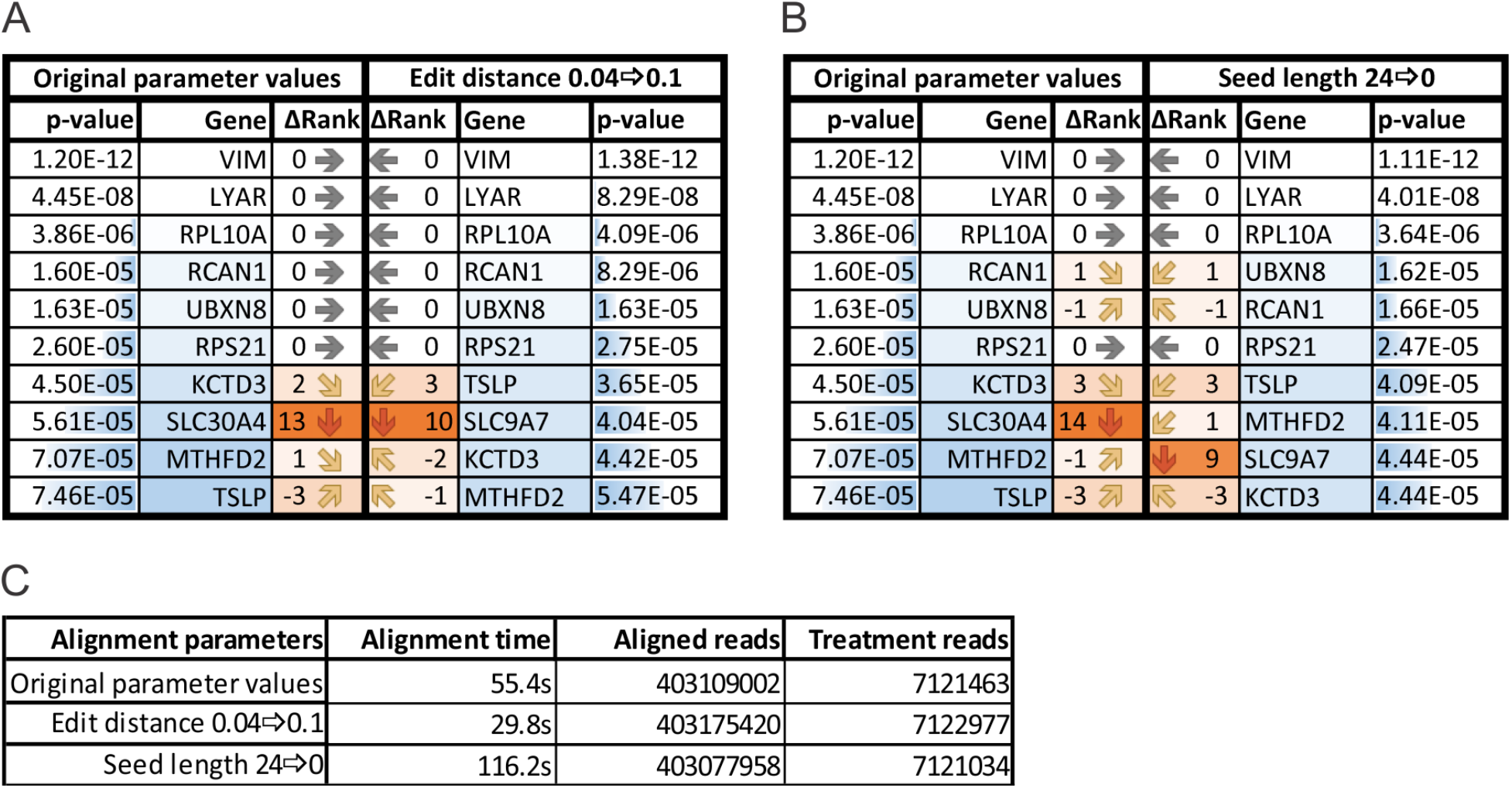
Changes in DEG lists after adjustment of alignment parameters. The effect on the top-10 DEG lists from cardiocytes treated with alendronate, increasing the maximum edit distance (1A), or not using a seed (1B) are shown. All genes are differentially expressed with a Benjamini-Hochberg False Discovery Rate of less than 0.1. The orange shading is proportional to the absolute value of the change in rank between the two lists. The arrow indicates where the gene is in the other list. In A, the gene SLC90A4 drops out of the top 10 with a drop of 13 ranks. Conversely, the gene SLC9A7, pops into the top 10 in the new list from position 18 in the original list. Similar changes are also seen in Table 1B when more accurate seedless alignment is used. Table C shows the alignment time for the entire dataset of 15 treatments on AWS excluding file transfer and function invocation times. The number of reads aligned for the entire dataset and the number of de-duped reads for alendronate treatment are also shown.

| Original parameter values |  |  | Edit distance 0.04⇒0.1 |  |  |
| --- | --- | --- | --- | --- | --- |
| p-value | Gene | ΔRank | ΔRank | Gene | p-value |
| 1.20E-12 | VIM | 0 → | ← 0 | VIM | 1.38E-12 |
| 4.45E-08 | LYAR | 0 → | ← 0 | LYAR | 8.29E-08 |
| 3.86E-06 | RPL10A | 0 → | ← 0 | RPL10A | 4.09E-06 |
| 1.60E-05 | RCAN1 | 0 → | ← 0 | RCAN1 | 8.29E-06 |
| 1.63E-05 | UBXN8 | 0 → | ← 0 | UBXN8 | 1.63E-05 |
| 2.60E-05 | RPS21 | 0 → | ← 0 | RPS21 | 2.75E-05 |
| 4.50E-05 | KCTD3 | 2 ↘ | ↗ 3 | TSLP | 3.65E-05 |
| 5.61E-05 | SLC30A4 | 13 ↓ | ↑ 10 | SLC9A7 | 4.04E-05 |
| 7.07E-05 | MTHFD2 | 1 ↘ | ↗ -2 | KCTD3 | 4.42E-05 |
| 7.46E-05 | TSLP | -3 ↗ | ↘ -1 | MTHFD2 | 5.47E-05 |

B
| Original parameter values |  |  | Seed length 24⇒0 |  |  |
| --- | --- | --- | --- | --- | --- |
| p-value | Gene | ΔRank | ΔRank | Gene | p-value |
| 1.20E-12 | VIM | 0 → | ← 0 | VIM | 1.11E-12 |
| 4.45E-08 | LYAR | 0 → | ← 0 | LYAR | 4.01E-08 |
| 3.86E-06 | RPL10A | 0 → | ← 0 | RPL10A | 3.64E-06 |
| 1.60E-05 | RCAN1 | 1 ↘ | ↗ 1 | UBXN8 | 1.62E-05 |
| 1.63E-05 | UBXN8 | -1 ↗ | ↘ -1 | RCAN1 | 1.66E-05 |
| 2.60E-05 | RPS21 | 0 → | ← 0 | RPS21 | 2.47E-05 |
| 4.50E-05 | KCTD3 | 3 ↘ | ↗ 3 | TSLP | 4.09E-05 |
| 5.61E-05 | SLC30A4 | 14 ↓ | ↑ 1 | MTHFD2 | 4.11E-05 |
| 7.07E-05 | MTHFD2 | -1 ↗ | ↘ 9 | SLC9A7 | 4.44E-05 |
| 7.46E-05 | TSLP | -3 ↗ | ↘ -3 | KCTD3 | 4.44E-05 |

**Table 1 Changes in DEG lists after adjustment of alignment parameters**
| Alignment parameters | Alignment time | Aligned reads | Treatment reads |
| --- | --- | --- | --- |
| Original parameter values | 55.4s | 403109002 | 7121463 |
| Edit distance 0.04⇒0.1 | 29.8s | 403175420 | 7122977 |
| Seed length 24⇒0 | 116.2s | 403077958 | 7121034 |

We have demonstrated how serverless cloud computing can be harnessed to provide on demand and affordable *“supercomputing”* made accessible with a graphical tool for RNA-seq analyses supporting interactive refinement of alignment parameters. We have shown how parameter tuning affects the ordering and composition of DEG lists even when bulk transcript counts are almost unaffected. DEG lists can be improved by avoiding compromises that compensate for slow execution. Our accelerated workflows enable scientists to explore parameter values to suit the application and data. Most importantly, the ability for non-cloud experts to interactively tune parameters in all steps of the analysis to iteratively examine the effect on DEG lists. This has implications for biological applications such as enabling more robust determination of diagnostic gene panels for personalized medicine.

## METHODS

### Basic workflow architecture

The workflow architecture is shown in **Figure S1**. The split and merge phases are performed on the client computer and a publish-subscribe mechanism is used to invoke the serverless instances to perform the alignment. Serverless functions are designed for small short-lived applications and accordingly have resource limitations. AWS Lambda functions are limited to 512 MB of disk space, 15 minutes maximum runtime, up to 3GB RAM, and 2 vCPUs^5^. Google functions are more limited with up to 2 GB RAM, part of which can be used as local disk space.

Due to the small and transient nature of the disk space allotted to a function, a storage bucket (AWS Simple Storage Service (S3) or Google bucket) is used to store input files that need to be transferred to a function, and output files that are generated by the function that need to be sent back to the client. The bucket acts as an intermediary layer for data transfer between the client and the serverless function instances, assuming the role of a distributed file system for a traditional supercomputer. Our design challenge was to modify the alignment process so it could execute within the limitations of serverless function instances while simultaneously reducing and mitigating the effect of the data transfers between the client, bucket, and serverless layers. This was accomplished by:

- Reduction of reference, data, and alignment file sizes
- Asynchronous non-blocking execution where the workflow proceeds with partial data
- Optimization of the client configuration
- Optimization of the serverless configuration

### Reduction of reference, data, and alignment file sizes

Data transferred to each serverless function instance includes the executables, input fastq files, and the human reference files. To reduce the size of reference files, we noted that BWA requires the reference sequence file, but only uses the name to infer filenames. To save space we created an empty dummy sequence file with the same name. Since we are aligning RNA sequences, we were also able to use the much smaller human exome reference, rather than the genomic reference. After the reference and executables are downloaded to the serverless instance, there is approximately 250 MB of space remaining for the input fastq and output alignment files. The original split step produced a single demultiplexed fastq file for each of the barcoded wells. We modified the executable to allow the user to produce multiple files no larger than a user-defined maximum size for each barcode. This process takes place on the client (e.g. laptop, local server, or cloud server) that then invokes the serverless functions. The maximum size of the input data also indirectly determines the number of serverless function instances that are invoked simultaneously. After experimenting with different file sizes on AWS, we determined that a data shard size of approximately 60 MB was near optimal, which enabled the use of 1752 serverless function instances to be invoked in parallel.

The most important file reduction was in the size of the alignment output files. BWA produces alignments using text-based SAM files that are approximately the same size as the input fastq files. Compression using gzip, or into BAM or the newer CRAM files can reduce the size by as much as 4-fold^15^. However, even the compressed files are relatively large. Also, compressing, and decompressing data files requires precious time especially when using the bzip algorithm which offsets the benefits of compression for reducing file transfer times. Instead, we developed a novel approach that reduced each alignment to a 64-bit hash value.

In the UMI-RNA-seq workflow, the Unique Molecular Identifier (UMI) is a random sequence attached to the read. Reads with the same UMI mapped to the same position are assumed to be amplification artifacts. Only one of these “duplicates” are counted. To detect these duplicates, we had previously implemented a one-to-one mapping, or perfect hash function, which combines the alignment position and the UMI to form a unique 64-bit value. Identical hash values are only possible if the reads map to the same position and have the same UMI, i.e. are amplification duplicates. To reduce output file sizes, the serverless instance pipes the output of BWA to a small executable (written in C++) that uses the same algorithm to convert the alignment and UMI into a hash value and then writes the hash to a binary file. The resulting binary files are 50x smaller than compressed SAM files, and do not require decompression to read. Once transferred to the bucket and downloaded to the client, the hash values are read in directly from the binary file and used by the merge executable to dedupe the reads. The alignment position is also recovered and used to obtain the transcript counts. A similar approach could also be used in non-UMI applications to encode just the alignment position.

### Asynchronous non-blocking execution

Our previous implementations proceeded sequentially through the split, align, and merge phases with each step beginning only after the prior step had completed. An important optimization was to refactor the splitting and upload of small files to the bucket layer to be asynchronous and nonblocking. Split files are now uploaded to buckets in parallel, as soon as they are written, instead of waiting for the entire split process to complete before transferring any files to the cloud. We also used the same optimization when downloading the starting fastq files to the client computer. Instead of waiting for all files to be downloaded before beginning the split phase, individual files are split as soon as they are downloaded. We did not apply this strategy to the alignment, download of alignments, and merge steps. For AWS using a fast cloud server, these steps were so fast that any additional benefit would not be worth the increase in complexity to the workflow. In the future, we may consider implementing asynchronous execution to improve performance for GCP functions and local server clients where these steps take longer to execute.

### Optimizing client configurations

The configuration of the client computer and the serverless function instances were also optimized. The longest steps in the workflow are the file splitting and upload steps which require good disk performance to read the fastq files, good single thread performance to process the files, and good network bandwidth to transfer the split files to the bucket layer. Most of the performance testing was done using different AWS instances spanning all families with local/resident SSDs, and we used equivalent GCP instances once we found an optimal AWS configuration.

#### Disk performance

The split executable speed is limited by the speed that it can read and write fastq files to and from the disk. AWS provides Elastic Block Storage (EBS) network drives, and local SSD’s for storage. The non-volatile memory (nVME) SSDs were faster than the fastest EBS drives, and performance was further enhanced by combining at least two drives in a RAID-0 array. We did not find further benefit from using additional disks in a RAID configuration.

#### Network performance

AWS provides different guaranteed throughput minimums. We did not obtain any benefit by increasing the network throughput beyond 25 GB/s. We also tried metal instances that do not use VMs but access the CPU directly. These instance types had significantly slower upload and download speeds to S3 buckets.

#### Single thread performance

The split executable is limited to file level parallelism – i.e. each pair of files is handled by a single thread. Thus, the optimal client should have multiple cores each with high single thread performance. For AWS, we used the z1 family of instances backed by the Intel Xeon Platinum 8151 12-core, 24-hyperthread CPU with a base clock speed of 3.4 GHz which bursts to 4.0 GHz CPUs to provide optimal single thread performance. In our initial testing, z1 instances were faster for the split step, than the c5 (Intel Xeon Platinum 8275CL CPU), and c6g (ARM-64 CPU) compute optimized instances. We did not test AMD based instances as they were not available with SSDs at the time of the benchmarking. Metal versions of these instances were slightly faster in execution of the split command due to the lack of virtualization overhead. However, this small advantage was outweighed by poorer upload/download speeds to and from S3 buckets.

#### Optimized cloud client

We found that the z1d.12xlarge AWS instance with 2 nVME SSDs in a RAID-0 configuration gave the best overall speed as a client for our workflow. The closest equivalent for GCP was the c2-standard-30 with 4 SSDs (the minimum available for this type of instance) and the premier network service tier. From the benchmarks in Table S1, we see that the GCP instance with more cores is slightly better for the merge operation which can theoretically use up to 96 threads, but slightly worse at the split function, likely due to poorer single thread performance.

### Optimizing serverless invocations

#### Invocation rate

The rate at which function instances can be invoked is a limiting factor in the speed of the alignment. For AWS Lambda, a burst limit allows 3000 function instances to be invoked almost instantaneously. Additional function instances are then provided at a rate of 500 per minute. However, AWS applies a default account limit on the number of concurrently running instances at 1000 which can be raised upon request. We also found that there was occasional undocumented throttling of the invocation rate when the number of concurrent instances exceeded one half of the limit. Therefore, for optimal performance on AWS, the number of the split fastq shards should not exceed 3000, and users should ask that the limit of concurrent instances be set to at least twice the number of shards. For Google functions, it is possible to set the number of concurrent instances to be unlimited. Using an unlimited number concurrent functions, it required approximately 3 minutes to invoke 1752 functions. It is possible that some of the function instances are re-used. Increasing the size of the shards to decrease the number of concurrent function invocations seemed to degrade performance, as the slightly shorter invocation time was offset by the longer execution time to align the larger shards.

#### Serverless invocation configuration

The pub-sub mechanism is used to start the execution of the serverless functions. The client publishes messages to a topic that the serverless function subscribes to. The client publishes messages containing the names of the files to be aligned. The serverless function reads these messages and spawns instances to retrieve the files from the bucket for alignment. On rare occasions, the alignment of some files is delayed, either from messages not being delivered promptly, or from some throttling of serverless invocations. Such a delay blocks the entire workflow, as all shards must be aligned before transcript counts can be obtained. To mitigate this problem, we monitored the start status of a serverless function by writing a start file to the bucket. We then would re-send the message from the client to the serverless function to process a missing shard if processing had not started within a reasonable time (set by the user).

#### Serverless instance configuration

AWS Lambda function instances provide up to two vCPUs and 3 GB available memory. The amount of available memory is configurable with additional memory being more expensive per second of execution. Using instances with the full 3 GB of memory allows both vCPUs to be used resulting in faster execution. We found that the additional per second cost of the extra RAM is almost completely offset by the savings in execution time. For Google functions, we were limited to 2GB memory shared between the disk space and available RAM. With only 2GB, there was not enough RAM to use both vCPUs simultaneously without swapping, so we only used 1 vCPU for alignment on Google functions. It may be beneficial in the case of Google functions to further reduce the size of the shards.

### Scripts for reproducing benchmarks

All Bash scripts used to benchmark our serverless RNA-seq workflow are available in the Github repository. The instructions for installing the scripts are given in the README. To facilitate installation on AWS, we also provide a script that begins from a basic Ubuntu image and installs all the executables, data, and scripts required to run the benchmarks.

### Installing and using graphical versions of the workflow

We provide a graphical user interface to orchestrate the serverless alignment as a completely interactive differential gene expression determination workflow. The graphical interface was built using the BioDepot-workflow-builder (Bwb). All modules in the workflow are containerized using Docker to ensure reproducibility and facilitate installation. Each module is represented by a graphical icon (widget). Double-clicking on a widget reveals tabbed windows for parameter entry and a display for monitoring the output of the widget. Right-clicking on the widget allows for customization of the interface of the widget. Dragging and dropping from one widget to another brings up dialogs for adding and deleting connections for data transfer and flow control. New modules from other workflows can also be added and connected by dragging-and-dropping. The workflows only require Docker to run, as everything, including Bwb is a container. Instructions on how to install Docker and use the workflows with the 46 GB of data in the case study or another dataset are given in a README in the repository

### Calculation of cost on AWS

The cost of the z1d.12xlarge client is based on the per hour on-demand Linux rate of ~$4.46 per hour. This rate includes the cost of the 2 SSDs attached to the instance. The running time was 6 min 16 seconds (0.1044 hours) for a cost of $0.47. The cost of the Lambda functions was calculated using the provided AWS calculator with the entries: 1752 executions, 3008 MB allocated memory, 39352 ms execution time and no free tier. The execution time is the average running time calculated over 10 sets of 1752 invocations. This results in an average cost for the Lambda functions of $3.38 for a total workflow cost of $3.85.

## SUPPLEMENTARY FIGURE AND TABLES

**Figure S1.**
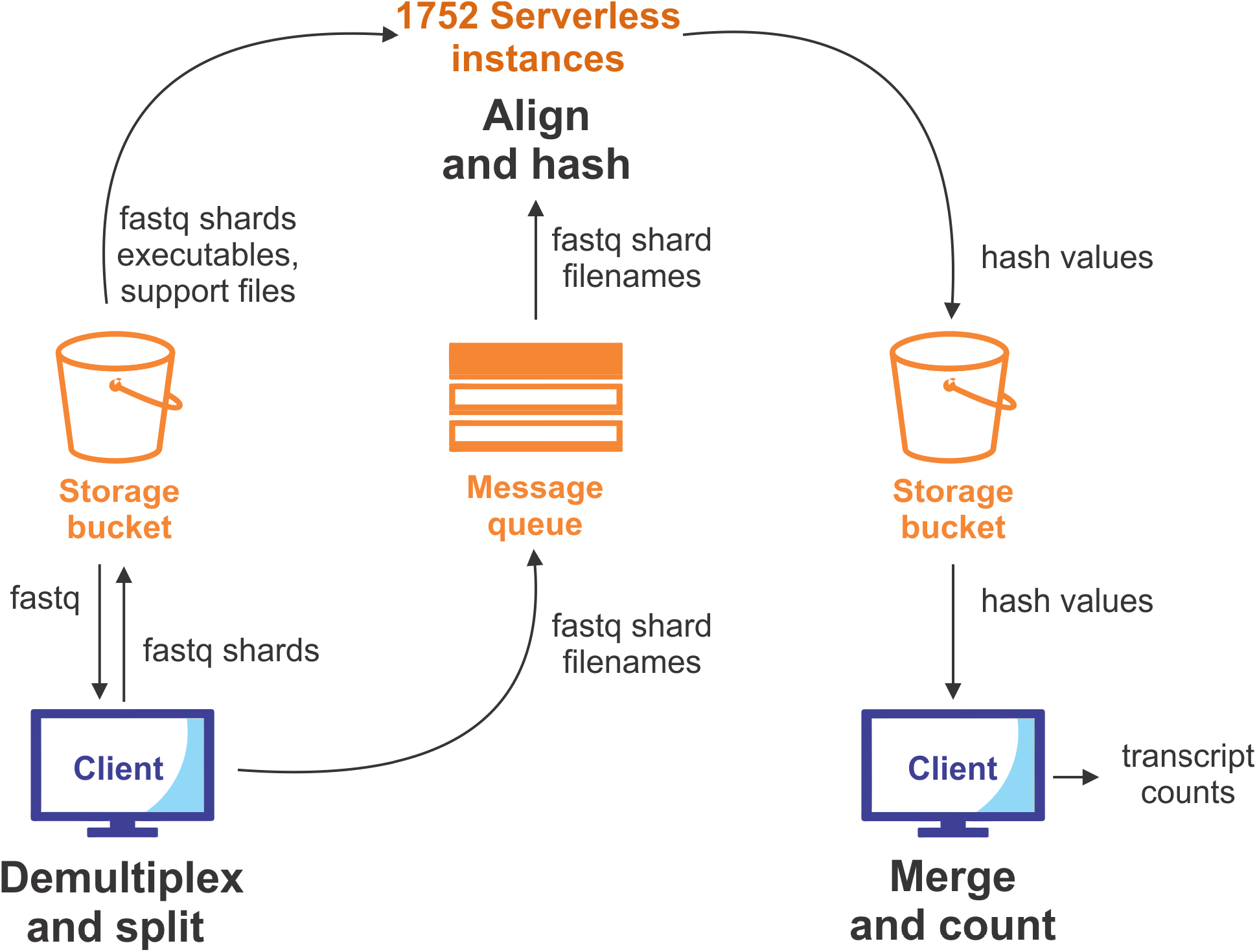
Data flow and architecture of serverless UMI-RNA-seq workflow. The workflow consists of 4 elements: *<u>a client</u>*-which executes the split and merge steps, *<u>serverless function instances</u>*-for alignment, *<u>a cloud storage bucket</u>*-for long term file storage and for temporary files in transit between the client and serverless instances, and *<u>a message queue</u>*-used to start the serverless instances and to receive status reports from the running instances. The client can be a laptop, local server, or cloud server. In the case of cloud clients, the starting fastq files are initially downloaded from a cloud storage bucket. The fastq files to be processed are demultiplexed and split into smaller pieces or shards. The small de-multiplexed fastq files are transferred to a storage bucket. When the split operation is complete, the client publishes messages containing the names of the sharded files to the message queue. Each function instance receives a message from the queue which specifies the fastq file it has been assigned to align. The function instance then downloads that shard from the storage bucket, along with all necessary executables and dependencies. The function instances run the alignment (with 2 threads for AWS, 1 thread for GCP) generating the 64-bit hash files as described in the Methods section. Hash files are then written to the storage bucket. The alignment is performed simultaneously for 1752 data shards using 1752 function instances. When all the hash files have been written to the storage bucket, the files are downloaded by the client and merged to obtain the transcript counts.

**Table S1.** Loss of gene from DEG list due to low counts. The top-10 DEG lists from cardiocytes treated with sunitinib and loperamide are shown. The data was processed using the original parameter values and using seedless alignment. The colorcoding scheme is the same as described for Table 1 with a blue gradient being used to denote the p-value, an orange gradient for the absolute value of the change in rank between the two lists. The arrows point to where the gene is in the other list. In addition, there is a logCPM (log Counts per Million reads) column with a green gradient bar that indicates the amount of expression found in the treated samples. For gene TNFRSF19, the gene counts were too low when processed with seedless alignment and edgeR filtered out the gene instead of calculating a logCPM. Consequently, it is not found in the lists of DEGs and may not be a robust gene to include in the top-10 list despite the highly significant p-value.

| Original parameter values |  |  |  | Seed length 24⇒0 |  |  |  |
| --- | --- | --- | --- | --- | --- | --- | --- |
| p-value | Gene | logCPM | ΔRank | ΔRank | logCPM | Gene | p-value |
| 1.01E-40 | HBB | 6.92 | 0 → | ← 0 | 6.92 | HBB | 7.13E-41 |
| 1.90E-07 | CDKN2B | 6.24 | 1 ↘ | ↘ 1 | 11.31 | VIM | 2.48E-07 |
| 2.63E-07 | VIM | 11.31 | -1 ↗ | ↗ -1 | 6.23 | CDKN2B | 2.97E-07 |
| 6.05E-07 | FABP4 | 4.65 | 0 → | ← 0 | 4.65 | FABP4 | 5.98E-07 |
| 7.30E-07 | LOC339524 | 0.83 | 0 → | ← 0 | 0.83 | LOC339524 | 6.97E-07 |
| 1.08E-06 | OXTR | 4.29 | 0 → | ← 0 | 4.29 | OXTR | 1.09E-06 |
| 1.59E-06 | TNFRSF19 | 0.69 | ## ↓ | ↘ 1 | 3.64 | PTPN13 | 1.95E-06 |
| 3.02E-06 | PTPN13 | 3.64 | -1 ↗ | ↘ 1 | 8.12 | CCL2 | 2.99E-06 |
| 3.19E-06 | CCL2 | 8.12 | -1 ↗ | ↘ 3 | 1.34 | CENPA | 9.76E-06 |
| 1.32E-05 | SYNM | 4.97 | 0 → | ← 0 | 4.97 | SYNM | 1.63E-05 |

**Table S2.** Alignment and total execution time for different UMI-RNA-seq implementations. The most performant implementation/configuration is highlighted in blue. See Table S3 for more details on the execution times. For the serverless implementations, alignment time is the execution time for the script that starts the serverless instances and does not terminate until all the alignment output has been uploaded to the storage bucket. Two values are shown for the total execution time. “Total” is the time including all file transfers, file splitting, and demultiplexing required only the first time the workflow is run. This also includes the time to transfer the 46 GB of compressed fastq files to the cloud-based clients. “Total (re-analysis)” is the time for subsequent re-analyses. The times for the original and optimized implementations are previously published median values from 3 runs on an AWS m4.4xlarge instance^5^. The serverless times are means from 5 runs using 1752 serverless instances for alignment (see Table S3 for all the run times). The local client was a 10 CPU core Xeon CPU E5-2640 v4 server. The cloud-based client on GCP was a c2-standard-30 instance with 60 virtual CPUs (vCPUs) and 4 solid state disks (SSDs). For AWS the client was a z1d.12x.large instance with 48 vCPUs and 2 SSDs. In both cases, the cloud-based SSDs were configured as a RAID-0 array.

| UMI RNA-seq implementation | Execution time (hour:minute:second) |  |  |
| --- | --- | --- | --- |
|  | Alignment | Total | Total (re-analysis) |
| Original | 19:38:09 | 29:25:27 | 23:54:22 |
| Optimized (16 threads) | 02:24:58 | 03:30:43 | 02:37:41 |
| Google Functions, local client | 00:05:13 | 00:21:13 | 00:06:59 |
| Google Functions, cloud client | 00:04:17 | 00:09:52 | 00:05:06 |
| AWS Lambda Functions, local client | 00:02:31 | 00:19:01 | 00:04:14 |
| **AWS Lambda Functions, cloud client** | **00:01:01** | **00:06:04** | **00:01:58** |

**Table S3.** Detailed stage execution times for different UMI-RNA-seq implementations. The individual run times used to calculate the mean run times in Table S2 are shown (h:m:s). Time for the split includes data transfer times of the fastq files to the cloud client, and data transfer times from the client to the cloud. Alignment times are the execution times for the script that starts the serverless instances and does not terminate until all the alignment output has been uploaded to the cloud bucket. Merge times include the time for transfer of aligned files to the client. Total1 is the time that would be required for reanalysis of the data, which would be the sum of the Align and Merge times. Total2 is the sum of all three phases, including the Split phase which is only required the first time the workflow is executed.

|  | AWS lambda functions: local client |  |  |  |  | AWS lambda functions cloud client |  |  |  |  |
| --- | --- | --- | --- | --- | --- | --- | --- | --- | --- | --- |
|  | Split | Align | Merge | Total1 | Total2 | Split | Align | Merge | Total1 | Total2 |
| Run 1 | 0:14:30 | 0:02:10 | 0:01:50 | 0:04:00 | 0:18:30 | 0:04:06 | 0:00:58 | 0:00:51 | 0:01:49 | 0:05:55 |
| Run 2 | 0:14:42 | 0:02:47 | 0:01:41 | 0:04:28 | 0:19:10 | 0:04:05 | 0:01:04 | 0:00:58 | 0:02:02 | 0:06:07 |
| Run 3 | 0:15:18 | 0:02:37 | 0:01:20 | 0:03:57 | 0:19:15 | 0:04:08 | 0:00:59 | 0:01:00 | 0:01:59 | 0:06:07 |
| Run 4 | 0:14:37 | 0:02:48 | 0:01:53 | 0:04:41 | 0:19:18 | 0:04:07 | 0:01:03 | 0:00:57 | 0:02:00 | 0:06:07 |
| Run 5 | 0:14:43 | 0:02:12 | 0:01:56 | 0:04:08 | 0:18:51 | 0:04:07 | 0:01:01 | 0:00:58 | 0:01:59 | 0:06:06 |
| Mean | 0:14:46 | 0:02:31 | 0:01:44 | 0:04:15 | 0:19:01 | 0:04:07 | 0:01:01 | 0:00:57 | 0:01:58 | 0:06:04 |
| Std dev | 0:00:19 | 0:00:19 | 0:00:15 | 0:00:19 | 0:00:20 | 0:00:01 | 0:00:03 | 0:00:03 | 0:00:05 | 0:00:05 |

**Table S3.** Detailed stage execution times for different UMI-RNA-seq implementations.
|  | Google functions: local client |  |  |  |  | Google functions: cloud client |  |  |  |  |
| --- | --- | --- | --- | --- | --- | --- | --- | --- | --- | --- |
|  | Split | Align | Merge | Total1 | Total2 | Split | Align | Merge | Total1 | Total2 |
| Run 1 | 0:13:52 | 0:05:35 | 0:01:46 | 0:07:21 | 0:21:13 | 0:04:50 | 0:04:30 | 0:00:47 | 0:05:17 | 0:10:07 |
| Run 2 | 0:14:18 | 0:04:35 | 0:01:46 | 0:06:21 | 0:20:39 | 0:04:41 | 0:04:19 | 0:00:51 | 0:05:10 | 0:09:51 |
| Run 3 | 0:14:18 | 0:05:51 | 0:01:45 | 0:07:36 | 0:21:54 | 0:04:42 | 0:04:29 | 0:00:49 | 0:05:18 | 0:10:00 |
| Run 4 | 0:14:25 | 0:05:33 | 0:01:47 | 0:07:20 | 0:21:45 | 0:04:44 | 0:04:09 | 0:00:49 | 0:04:58 | 0:09:42 |
| Run 5 | 0:14:15 | 0:04:32 | 0:01:46 | 0:06:18 | 0:20:33 | 0:04:52 | 0:03:59 | 0:00:50 | 0:04:49 | 0:09:41 |
| Mean | 0:14:14 | 0:05:13 | 0:01:46 | 0:06:59 | 0:21:13 | 0:04:46 | 0:04:17 | 0:00:49 | 0:05:06 | 0:09:52 |
| Std dev | 0:00:13 | 0:00:37 | 0:00:01 | 0:00:37 | 0:00:37 | 0:00:05 | 0:00:13 | 0:00:01 | 0:00:13 | 0:00:11 |

## ACKNOWLEDGEMENTS

L.H.H, W.L., X.N. and K.Y.Y. are supported by National Institutes of Health (NIH) grant R01GM126019. L.H.H and X.N. are also supported by University of Washington CoMotion Step Fund and Innovation Gap Fund. WL is also supported by NSF grant OAC-1849970. This project has been funded in whole or in part with Federal funds from the National Cancer Institute, National Institutes of Health, Department of Health and Human Services, under Contract No. 75N91020C00009.

We would also like to thank Google Cloud Platform (K.Y.Y.), Amazon Web Services (L.H.H., W.L., and K.Y.Y.) for computing resources. We would like to thank Dimitar Kumanov for contributions to an early version of this work.

